# Silencing *KRIT1* Partially Reverses the Effects of Disturbed Flow on the Endothelial Cell Transcriptome

**DOI:** 10.1101/2025.03.12.642862

**Authors:** Amelia Meecham, Sara McCurdy, Eduardo Frias-Anaya, Wenqing Li, Helios Gallego-Gutierrez, Phu Ngyuen, Julie Yi-Shuan Li, Shu Chien, John Y.-J. Shyy, Mark H. Ginsberg, Miguel A. Lopez-Ramirez

## Abstract

**Background:** Endothelial cells respond to forces generated by laminar blood flow with changes in vasodilation, anticoagulant, fibrinolytic, or anti-inflammatory functions which preserve vessel patency. These responses to flow sheer stress are primarily mediated by the modulation of transcription factors Krüppel-like factors 2 and 4 (KLF2 and KLF4). Notably, disturbed flow patterns, which are found in vascular areas predisposed to atherosclerosis, significantly reduce the endothelial expression of KLF2 and KLF4, resulting in changes in the transcriptome that exacerbate inflammation and thrombosis. The endothelial CCM complex, comprising KRIT1, CCM2, and CCM3, suppresses the expression of KLF2 and KLF4. Loss of function of the CCM complex has recently been suggested to protect from coronary atherosclerosis in humans. We thus hypothesized that silencing of *KRIT1*, the central scaffold of the CCM complex, can normalize the atherogenic effects of disturbed flow on the human endothelial transcriptome

**Methods:** Bulk RNA sequencing (RNA-seq) was conducted on human umbilical vein endothelial cells (HUVECs) after the expression of KRIT1 was silenced using specific siRNAs. The endothelial cells were exposed to three different conditions for 24 hours: pulsatile shear stress (laminar flow), oscillatory shear stress (disturbed flow), and static conditions (no flow).

**Results:** We found that silencing *KRIT1* expression in HUVECs restored the expression of the transcription factors KLF2 and KLF4 under oscillatory shear stress. This treatment resulted in a transcriptomic profile similar to that of endothelial cells under pulsatile shear stress. These findings suggest that inhibition of the CCM complex in endothelium plays a vasoprotective role by reactivating a protective gene program to help endothelial cells resist disturbed blood flow.

**Conclusions:** Targeting CCM genes can activate well-known vasoprotective gene programs that enhance endothelial resilience to inflammation, hypoxia, and angiogenesis under disturbed flow conditions, providing a novel pathway for preventing atherosclerosis.

**Graphic abstract:** 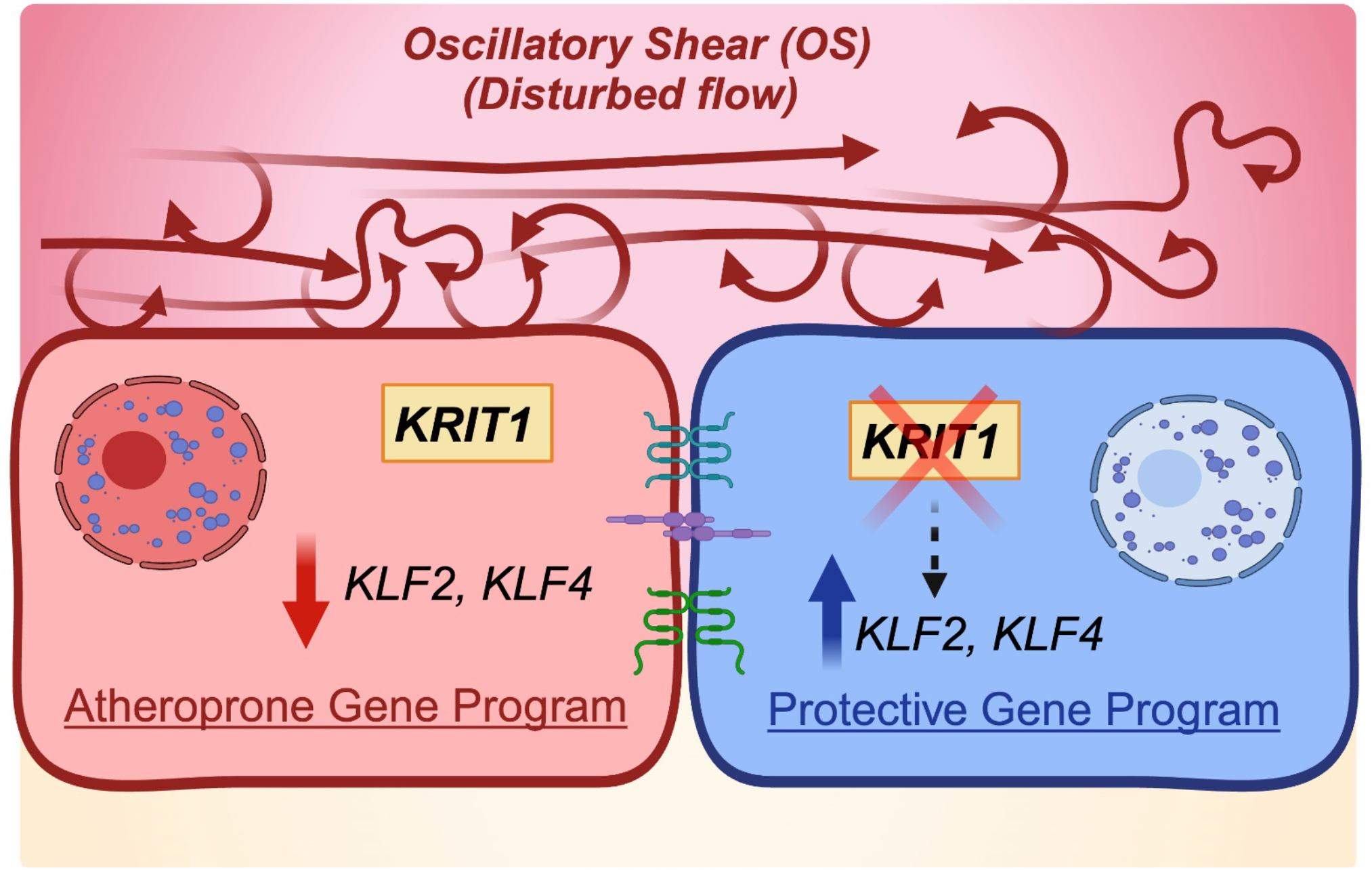

Endothelial cells respond to blood flow by expressing transcription factors such as Krüppel-Like Factors 2 and 4 (KLF2 and KLF4). Under oscillatory shear stress (OS) conditions, characterized by disturbed flow, KLF2 and KLF4 are suppressed in endothelial cells, which is linked to an atheroprone gene program that includes increased inflammation and risk of atherosclerosis (unhealthy endothelium). Our work shows that silencing of endothelial KRIT1 under OS increases expression of *KLF2* and *KLF4* mRNA levels as well as the expression of a protective gene program associated with resilience to inflammation, hypoxia, and angiogenesis, similar to those observed under pulsatile shear stress conditions (healthy endothelium).

## Introduction

Inflammation in cardiovascular diseases is a significant and common clinical issue and morbidity is often due to thrombosis and endothelial dysfunction^1^. Healthy, quiescent vascular endothelial cells have anticoagulant, fibrinolytic, and anti-inflammatory properties^2,3^. Flow patterns are known to have specific effects on vascular endothelial gene expression, and have been the focus of numerous studies, highlighting the importance of specific genetic programs in maintaining endothelial homeostasis ^4-6^. The vasoprotective effects of laminar blood flow are primarily driven by the upregulation of the transcription factors KLF2 and KLF4 (Krüppel-like factors 2 and 4)^7,8^. These factors significantly enhance the expression of genes encoding anticoagulants, such as *THBD*, which encodes thrombomodulin (TM), and vasodilators, like *NOS3*, which encodes endothelial nitric oxide synthase (eNOS). KLF2 and KLF4 suppress the expression of genes that inhibit angiogenesis, including *THBS1*, which encodes thrombospondin-1 (TSP1), as well as pro-inflammatory genes driven by NF-κB, such as vascular adhesion molecules *VCAM1* and *ICAM1*^7,9-11^. Moreover, KLF2 and KLF4 are regulators of hypoxia by promoting degradation of Hypoxia-inducible factor 1 (HIF-1)^12^. These vasoprotective effects of laminar blood flow have inspired new therapeutic strategies centered around enhancing the expression of endothelial KLF2 and KLF4^7,13,14^, to reduce cardiovascular morbidity^15,16^.

Recent *in vitro* studies have documented the gene expression programs in cultured human umbilical vein endothelial cells (HUVECs) and human aortic endothelial cells (HAECs) exposed to different patterns of shear stress^17^. Pulsatile shear stress (laminar flow) promotes an atheroprotective endothelial phenotype (e.g., by increasing eNOS, Cholesterol-25-hydroxylase, Ch25h, Liver X receptor, LxR, Cycle-dependent kinase 2, CDK2), while oscillatory shear stress (disturbed flow) is linked to an atheroprone phenotype (e.g., by increasing NF-κB signaling and oxidative stress)^6,17-21^. Using these culture systems, key mechanotransduction pathways vital for endothelial function and homeostasis have been identified^17-19,22-25^. For instance, oscillatory shear significantly downregulates KLF2 and KLF4 in endothelial cells, leading to dysregulation of many downstream vasoprotective gene programs^19^.

A recent study utilizing Perturb-seq, a method combining CRISPR-based gene silencing with single-cell RNA sequencing, investigated the effects of individually silencing 300 genes in aortic endothelial cells. These genes were either associated with or located near SNPs linked to an increased risk of coronary artery disease (CAD). The analysis showed that the cerebral cavernous malformation (CCM) signaling pathway is connected to CAD risk genes and influences atheroprotective processes in endothelial cells^26^.

In this study, we tested the hypothesis that silencing *KRIT1* would promote vasoprotective transcriptional responses in endothelial cells exposed to disturbed flow^6^. We performed bulk RNA sequencing (RNAseq) on HUVEC treated with *KRIT1* siRNA and exposed to pulsatile, oscillatory, or no flow for 24 hours. Our findings indicate that targeting the endothelial CCM complex may help restore the expression of vasoprotective gene programs downregulated by oscillatory shear stress. By mimicking a pulsatile shear stress gene program, this approach may help negatively regulate inflammation, hypoxia, and angiogenesis in mature vascular endothelial cells, ultimately protecting against atherothrombosis.

## Material and methods

### Cell culture and transfection with siRNA

HUVECs purchased from Lonza (C2519A) were cultured according to the manufacturer’s protocol and recommended growth medium (CC-3162). Experiments described were performed using cells between passages 3 and 6. HUVECs were maintained by passaging 1:3 and grown to confluence in fibronectin-coated T-75 flasks using 2μg/mL human fibronectin (Sigma, F2006) dissolved in sterile PBS and incubated for 1 hour at 37°C. For in vitro experiments, cells were transfected with 75nM SMARTpool siRNA against KRIT1 (Dharmacon, M-003825–01) or non-targeting control siRNA (Dharmacon, D-001206–13). Transfection mix was prepared in OptiMEM (Gibco) using lipofectamine3000 (Invitrogen), which was added dropwise to cells plated in complete growth medium. Following overnight (12-hour) incubation, cells were given fresh growth medium for 24 hours before replating onto fibronectin-coated glass slides for flow chamber experiments.

### Flow chamber experiments

HUVECs transfected with siRNA for 48 hours were replated using 0.05% Trypsin/EDTA (Thermo Fisher, 25300054) on 38 mm × 76 mm glass plates at 95% confluence. A subset of the glass plates was then assembled into a parallel-plate flow channel as previously described^27^ for each condition: siSCR+PS, siKRIT1+PS, siSCR+OS, and siKRIT1+OS. The flow system was maintained at 37°C and ventilated with 95% humidified air with 5% CO_2_. HUVECs were exposed to pulsatile flow (PS) (12 ± 4 dyn/cm2, 1 Hz) or oscillatory flow (OS) (1 ± 4 dyn/cm2, 1 Hz) for 24 hours as described previously^28^. Additional control plates of siSCR- and siKRIT1-transfected HUVECs were subject to static conditions (no flow) and maintained in the same culture medium and conditions described above in an adjacent incubator.

### RNA isolation

Glass slides of HUVECs were removed from the flow chamber and cells were directly lysed in 400µl of TRIzol (Invitrogen, 15596026) per slide and stored in RNase-free microcentrifuge tubes, which were transferred to dry ice for storage until all samples were collected. Total RNA was extracted using the RNeasy mini kit (Qiagen, 74104) according to manufacturer’s directions. RNA quality and quantity was measured on a NanoDrop spectrophotometer (Thermo Fisher) before cDNA synthesis using the qScript cDNA SuperMix First-Strand cDNA Synthesis (Quantabio, 95048-100). Silencing of *KRIT1* transcript and expression of known downstream genes was verified by quantitative real-time PCR (qRT-PCR) before samples were submitted to the UCSD IGM Genomics Center for RNA integrity analysis and bulk RNA sequencing.

### RNA-sequencing (RNA-seq) analysis

RNA quality was confirmed using an Agilent Tapestation 4200, and high-quality RNA (RIN > 8.0) was used for RNA-seq library preparation with the Illumina® Stranded mRNA Prep kit (Illumina, San Diego, CA). RNA libraries were multiplexed and sequenced with 100-bp paired reads (PE100) on an Illumina NovaSeq 6000 platform with a target depth of 30 million reads per sample. Samples were demultiplexed using bcl2fastq conversion software (Illumina, San Diego, CA).

Sequencing analysis was conducted in the R environment, following quality assurance of the data using FastQC. Reads were aligned to the human genome (GRCh38) using RNAStar^29^, and read counts were obtained using featureCounts^30^. Differential expression analysis and normalization of counts were performed using DESeq2^31^. RNA-seq data from this study are accessible in the Gene Expression Omnibus (GEO) database (GSE288811, http://www.ncbi.nlm.nih.gov/geo). The corresponding authors can be contacted for requests regarding public and non-public data supporting this study’s findings.

### Statistical analysis

Statistical analyses were performed using GraphPad Prism software. Data are presented as mean ± standard error of the mean (SEM) across multiple biological experiments, with the number of independent replicates specified for each experiment. For comparisons between groups, multiple unpaired t-tests with the Holm-Sidak method were used to adjust for multiple testing, with a significance threshold set at p < 0.05.

## Abbreviations and Acronyms

CCMs: Cerebral Cavernous Malformations
KRIT1: Krev1 interaction trapped gene 1
HEG1: Heart of glass 1
CCM2: Malcavernin
RNA-seq: RNA sequencing
KLF2: Krüppel-like factors 2
KLF4: Krüppel-like factors 4
TM: Thrombomodulin
eNOS: Endothelial nitric oxide synthase
TSP1: Thrombospondin-1
VCAM1: Vascular cell adhesion molecule 1
ICAM1: Intercellular adhesion molecule 1
NF-κB: Nuclear Factor kappa B
HUVEC: Human umbilical vein endothelial cells
HAEC: Human aortic endothelial cells
CAD: Coronary artery disease

## Author contributions

A.M., S.M. designed and performed the experiments, analyzed and interpreted the data, generated the figures, and wrote the manuscript. E.F-A., W.L., and H.G-G. analyzed and interpreted the data, generated the figures, and helped write the manuscript. P.N., J.L., S.C., JS., performed shear stress experiments. M.H.G., M.A.L-R., conceived the project, designed the overall study, analyzed and interpreted data, and wrote the manuscript.

## Results

### Transcriptomic responses of KRIT1-targeted HUVECs when subjected to oscillatory and pulsatile shear stress or static conditions

To evaluate whether silencing of *KRIT1* can enhance the expression of the transcription factors KLF2 and KLF4 in endothelial cells experiencing different patterns of sheer stress, we subjected *KRIT1* siRNA-treated HUVECs to pulsatile sheer (PS), oscillatory shear (OS), or no flow (static) for 24 hours (Figure 1A). We observed that silencing *KRIT1* (siKRIT1) led to increased levels of the transcription factors *KLF2* and *KLF4*, as well as the KLF target gene, *NOS3* across all conditions, as measured by RT-qPCR (Figure 1B). Conversely, downregulation of these important atheroprotective genes was observed in siSCR control HUVECs subjected to OS compared with PS conditions (Figure 1B). These data agree with previous studies reporting the same regulation of these flow-responsive genes under laminar (PS) or disturbed (OS) flow, both *in vitro* and *in vivo*^17^. Interestingly, targeting *KRIT1* expression (siKRIT1) in HUVECs exposed to OS significantly restored expression of *KLF2* and *KLF4* and *NOS3* mRNA to levels observed in siSCR control cells under PS (Figure 1B).

**Figure 1.**
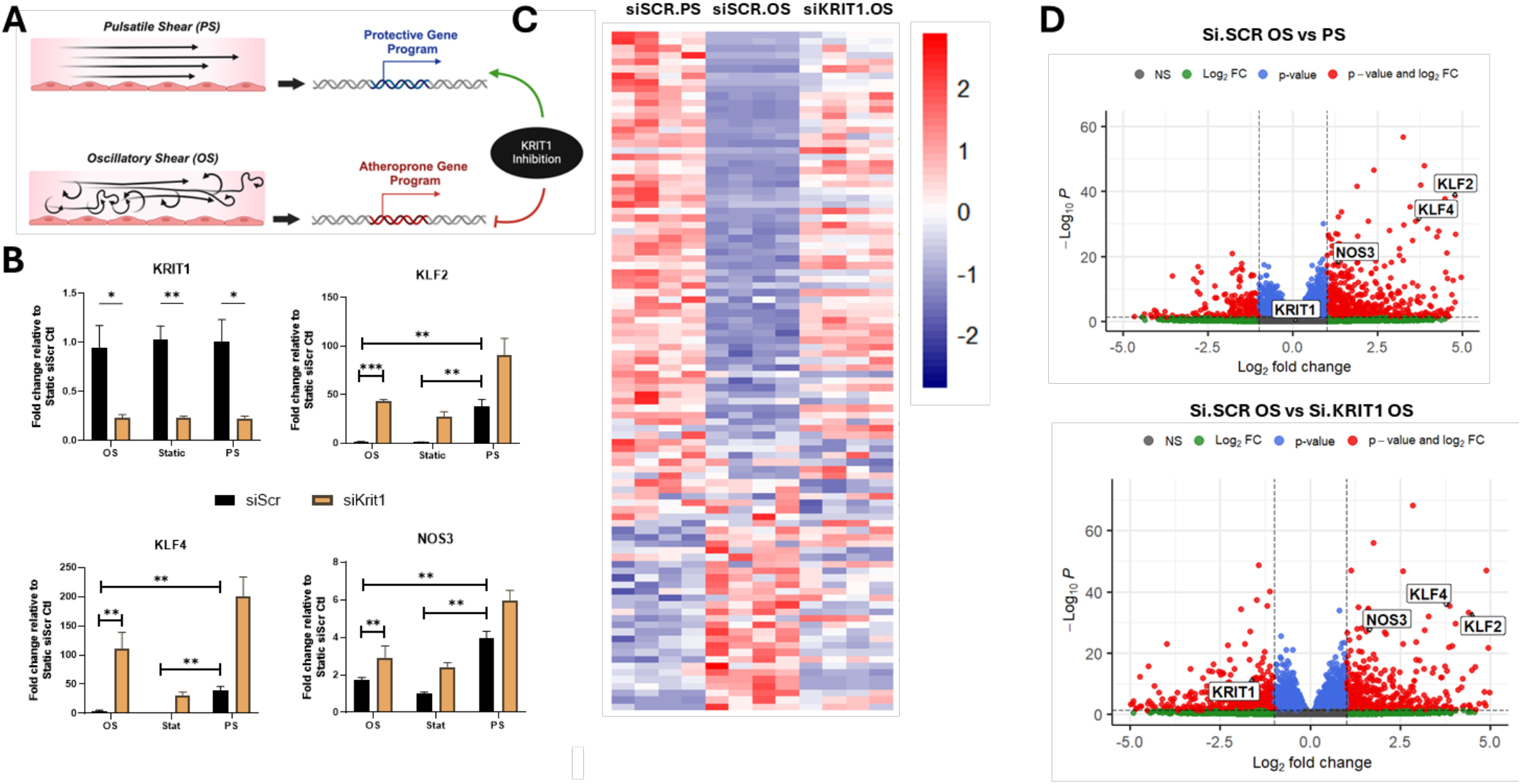
Transcriptomic responses of *KRIT1*-targeted HUVECs subjected to different flow patterns. **A)** Schematic depicting the relationship between flow type and endothelial response, with KRIT1-targeting leading to a transcriptomic profile that is most similar to that seen in cells experiencing vasoprotective pulsatile flow. **B)** RT-qPCR analysis of key vasoprotective genes (*KLF2, KLF4, NOS3)* and confirmation of *KRIT1* silencing in scrambled siRNA (siSCR)- or *KRIT1* siRNA (siKRIT1)-treated HUVECs exposed to no flow (static), pulsatile shear (PS), or oscillatory shear (OS) for 24 hours. (n=4, *p<0.05, **p<0.001, ***p<0.0001). **C)** Heatmap of top 100 differentially expressed genes comparing PS to OS flow under control conditions following bulk RNAseq of the samples described in panel A (n=4) **D)** Volcano plots from bulk RNASeq. **Top:** comparison of PS and OS flow under control conditions highlighting higher expression of key vasoprotective genes (KLF2, KLF4, NOS3) under PS. **Bottom:** comparison of transcriptome in control and KRIT1 knockdown cells under OS, highlighting the successful knockdown of *KRIT1* and upregulation of KLF2, KLF4, and NOS3 in siKRIT1-treated cells relative to siSCR control. Red points represent genes that are >2 fold changed and p<0.05 green points indicate a genes that are >2 Fold changed but p>0.05. Blue points indicate genes that are <2fold changed. Grey points indicate non-significantly changed genes,

To better understand the transcriptomic profile of changes due to KRIT1-targeting, we performed bulk RNA sequencing (RNAseq) on the groups of HUVECs as described above; *KRIT1* knockdown (siKRIT1) vs control (siSCR) subjected to PS or OS. Significant differentially expressed genes were determined by Deseq adjusted p values (p<0.05) and at least a log fold change of 1. We observed that KRIT1-targeted HUVECs under OS exhibited a transcriptomic profile that closely resembled that of HUVECs treated with control siRNA under PS (Figure 1C). Notably, consistent with RT-qPCR results, we observed that *KRIT1-*targeted HUVECs under OS increased expression of transcription factors *KLF2* and *KLF4* (Figure 1D).

### Silencing *KRIT1* in endothelial cells experiencing disturbed flow activates protective gene programs known to enhance resistance to inflammation, hypoxia, and angiogenesis

To investigate the transcriptional effects of KRIT1-targeted HUVECs, we conducted pathway analysis using the differentially expressed genes (DEGs) identified comparing siKRIT1 HUVECs to the siSCR HUVECs. A total of 59 pathways were found to be significantly enriched under OS, including those related to cell-matrix adhesion, response to shear stress, integrin-mediated signaling, response to hypoxia, and regulation of angiogenesis (Figure 2A, Supplemental Feigure 1). For inflammation pathway genes, the expression profile of siKRIT1 cells exposed to OS showed notable similarities to those of siSCR HUVEC under PS, including the elevation of CMKLR1, IL17RE, TBXA2R and downregulation of JUN, SPHK1, P2RX7 and C2CD4B (Figure 2B). In hypoxia pathways, the gene expression patterns in siKRIT1 HUVEC under OS again aligned with those seen in siSCR HUVEC under PS, including the elevation of AQP1, ADRB2, HIF3A, further supporting a potential protective transcriptional response against OS-induced hypoxic stress. We also observed that with angiogenesis pathway genes, siKRIT1 HUVEC under OS also showed expression changes similar to those seen in control cells under PS. Notably, CYP1B1, APLNR, RAMP2, and SEMA5A were increased. These results indicate that KRIT1 silencing in HUVEC exposed to disturbed flow leads to a change in transcriptome associated with vascular protection.

**Figure 2.**
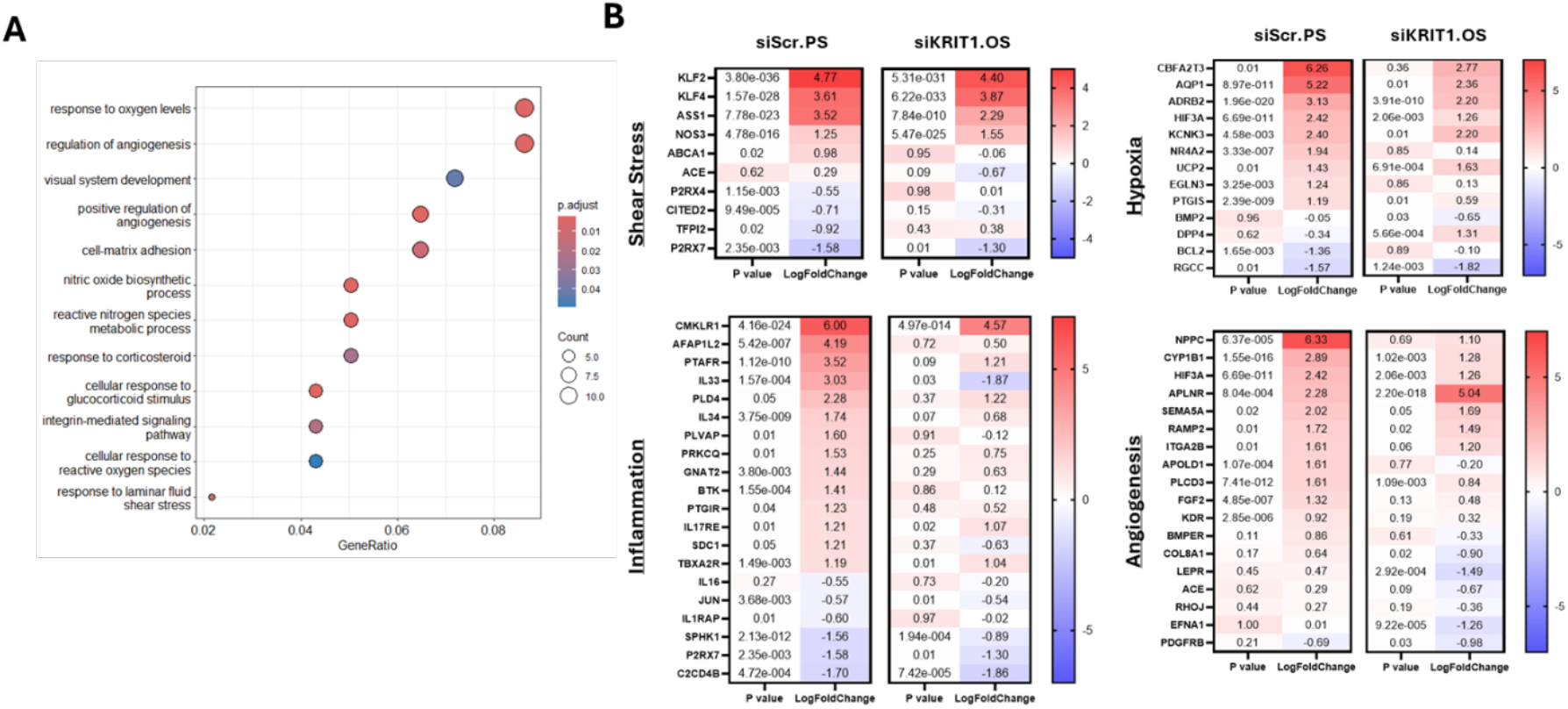
Biological pathway analysis of KRIT1-targeted HUVECs under disturbed flow conditions. **A)** Dot plot highlighting a selection of the significantly enriched biological pathways based on the DEGs in response to OS in both control and *KRIT1*-knockout cells. Pathway analysis was performed using the ‘clusterProfiler’ package in R. **B)** Heatmaps of selected genes from four of the significantly changed pathways. P value and LogFoldChange generated by Deseq2 package. (n=4)

## Discussion

We demonstrate that targeting endothelial KRIT1 initiates a protective response in endothelial cells exposed to disturbed flow patterns, which helps prevent the onset of an atheroprone gene program associated with endothelial dysfunction. In our study, we utilized flow chamber experiments to expose endothelial cells to various conditions, including oscillatory shear stress, pulsatile shear stress, and static conditions. Our analysis indicated that targeting KRIT1 expression in HUVEC under these three conditions resulted in a significant increase in the levels of the transcription factors KLF2 and KLF4, as well as in protective gene programs that enhance resilience to inflammation, hypoxia, and angiogenesis in the face of disturbed flow conditions.

It is well-established that vascular endothelial cells undergo transcriptional changes in response to specific flow patterns. Exposure to oscillatory shear stress *in vitro* triggers transcriptomic responses mimicking those observed in the endothelium of athero-prone aortic regions. Likewise, endothelial cells subjected to pulsatile shear stress *in vitro* exhibit gene expression patterns resembling those found in vascular regions with laminar flow, which are known to be resistant to atherosclerotic plaque formation ^32,33^. The flow-responsive transcription factors KLF2 and KLF4 play an important role in restricting endothelial dysfunction, inflammation, thrombosis, and angiogenesis in response to disturbed flow^34,35^.

The link between the CCM complex and cardiovascular disease risk is the focus of a recent study by Schnitzler *et al*, responsible for identifying 43 coronary artery disease GWAS signals that all converged on the CCM complex pathway. The authors went on to show that CRISPR knockdown of *CCM2*, the main binding partner of KRIT1, in endothelial cells leads to activation of anti-inflammatory, anti-thrombotic, barrier-promoting genetic programs^26^. While KRIT1 is considered the central scaffold of the CCM protein complex, it also has a transmembrane binding partner, heart of glass 1 (HEG1), which is responsible for anchoring KRIT1 at endothelial junctions^36^.

Our own recent work has shown that the interaction between KRIT1 and HEG1 can be pharmacologically inhibited, resulting in increased endothelial expression of KLF2, KLF4, and eNOS ^15^, establishing the endothelial CCM complex as a potentially druggable target, with small molecule inhibitors a potential therapeutic option for prevention of endothelial dysfunction in vascular disease.

Pharmacologically inhibiting the endothelial KRIT1-HEG1 protein interaction with a compound denoted HKi2^15^ under static conditions increases the expression of KLF4 and KLF2 and mimics the vasoprotective transcriptional effects of laminar blood flow^15,37^. Here, we extend those studies in two ways: 1) we show that genetic silencing of *KRIT1* triggers similar transcriptional programs to those initiated by HKi2 and 2) we show that under conditions of disturbed flow, known to trigger transcriptional changes associated with predisposition to atherosclerosis, that silencing *KRIT1* leads to a transcriptional program more like that observed in atheroprotective laminar flow.

These findings align with a recent study that showed that blood flow can drive the release of endothelial KRIT1 from the CCM complex and trigger the activation of a MEKK3-MEK5-ERK5-MEF2 pathway^38^ that increases KLF2 and KLF4 expression^38^. Furthermore, the HEG1 protein migrates to the downstream side of the cells and is secreted into the medium alongside KRIT1^38^. Depleting KRIT1-HEG1 from endothelial cells triggers the activation of the MEKK3-MEK5-ERK5-MEF2 pathway, which is a well-known regulator of KLF2 and KLF4 expression^7,16,39^. Future research should focus on gaining a deeper understanding of the biology of the endothelial CCM protein complex. This understanding is essential to clarify the molecular mechanisms that differentiate between beneficial and protective processes and those that result in harmful dysfunction in endothelial cells associated with CCM disease^40-42^.

Taken together, these studies, along with our findings, strengthen the emerging paradigm that KRIT1 regulates endothelial adaptation to disturbed flow through transcriptional regulation of vasoprotective pathways. Furthermore, they highlight the potential of CCM complex proteins as therapeutic targets for vascular disease.

